# Up-regulation of autophagy by low concentration of salicylic acid delays methyl jasmonate-induced leaf senescence

**DOI:** 10.1101/814608

**Authors:** Runzhu Yin, Jingfang Yu, Yingbin Ji, Jian Liu, Lixin Cheng, Jun Zhou

**Author notes:** Correspondence: Lixin Cheng,; Jun Zhou.

## Abstract

Crosstalk between salicylic acid (SA) and jasmonic acid (JA) signaling plays an important role in molecular regulation of plant senescence. Our previous works found that SA could delay methyl jasmonate (MeJA)-induced leaf senescence in a concentration-dependent manner. Here, the effect of low concentration of SA (LCSA) application on MeJA-induced leaf senescence was further assessed. High-throughput sequencing (RNA-Seq) results showed that LCSA did not have dominant effects on the genetic regulatory pathways of basal metabolism like nitrogen metabolism, photosynthesis and glycolysis. The ClusterONE was applied to identify discrete gene modules based on protein-protein interaction (PPI) network. Interestingly, an autophagy-related (ATG) module was identified in the differentially expressed genes (DEGs) that exclusively induced by MeJA together with LCSA. RT-qPCR confirmed that the expression of most of the determined ATG genes were upregulated by LCSA. Remarkably, in contrast to wild type (Col-0), LCSA cannot alleviate the leaf yellowing phenotype in autophagy defective mutants (*atg5-1* and *atg7-2*) upon MeJA treatment. Confocal and western blot results showed that LCSA increased the number of autophagic bodies and autophagic flux during MeJA-induced leaf senescence. Collectively, our work revealed up-regulation of autophagy by LCSA as a key regulator to alleviate MeJA-induced leaf senescence.

## INTRODUCTION

Senescence in green plants is a complex and orderly regulated process that is crucial for transiting from nutrient assimilation to nutrient remobilization (Masclaux et al., 2000; Quirino et al., 2000; Lim et al., 2003; Yoshida, 2003; Schippers, 2015). During senescence, the most visible characteristic is leaf yellowing, which is the consequence of a succession of changes in cellular physiology including chlorophyll degradation and photosynthetic activity reduction (Lim et al., 2003; Yoshida, 2003). Chloroplast as an early senescence signaling response organelle, its dismantling plays an important role in the major nitrogen source recycling and remobilization (Avila-Ospina et al., 2014). The progression of leaf senescence can be prematurely induced by multiple environmental and endogenous factors, such as temperature, light, humidity and phytohormones (Lim et al., 2007). Hormone signaling pathways play roles at all the stages of leaf senescence, including the initiation, progression, and the terminal phases of senescence (Lim et al., 2007). Recent progresses show that senescence can be coordinately regulated by several phytohormones like cytokinins, ethylene, abscisic acid, salicylic acid (SA), and jasmonic acid (JA) (Gan and Amasino, 1995; van der Graaff et al., 2006; Hung and Kao, 2004; He et al., 2002; Morris et al., 2000). However, the detailed molecular mechanisms for these phytohormone signals in plant senescence remain poorly understood.

JA has been known as a key plant hormone for promoting senescence, based on the findings that exogenously applied methyl jasmonate (MeJA, methyl ester of JA) leads to a rapid loss of chlorophyll content and accompany with reduction of photochemical efficiency (Yue et al., 2012; Ji et al., 2016). Studies with JA-insensitive mutant *coronatine insensitive 1* (*coi1*) that exhibited defective senescence response to MeJA treatment (He et al., 2002), supporting the notion that JA signaling pathway is crucial for leaf senescence. Some other evidences indicate that SA is also involved in plant senescence (Morris et al., 2000; Chai et al., 2014). The concentration of endogenous SA increases to upregulate several senescence-associated genes during leaf senescence (Morris et al., 2000; Yoshimoto et al., 2009). However, such genetic regulatory mechanisms are abolished in plants defective in the SA signaling or biosynthetic pathway (*npr1* and *pad4* mutants, and *NahG* transgenic plants) (Morris et al., 2000). Crosstalk between MeJA and SA has been broadly documented in plant defense response, which commonly manifests as a reciprocal antagonism pattern (Thaler et al., 2012). Evidence suggests that antagonistic interactions between SA and MeJA modulate the expression of a senescence-specific transcription factor WRKY53, showing induced by SA, but repressed by MeJA (Miao and Zentgraf, 2007). Overall, mechanisms determining the specificity and coordination between SA and JA still need to be further explored.

Most of phytohormones have both stimulatory and inhibitory effects on the growth and metabolism of higher plants in a dose dependent manner. It seems that SA functions in the same way on the physiological and biochemical processes of plants (Ji et al., 2016; Pasternak et al., 2019). Low-concentration SA (hereafter as LCSA) at below 50 micromole (µM) promotes adventitious roots and altered architecture of the root apical meristem, whereas high-concentration SA (greater than 50 µM) inhibits root growth (Pasternak et al., 2019). Interestingly, we previously demonstrated that MeJA-induced leaf senescence could be delayed by LCSA (1-50 μM), but accelerated when the concentration higher than 100 µM (Ji et al., 2016). Our other related works have verified such high dose of SA greatly activates NPR1 (nonexpressor of pathogenesis-related genes 1) translocation into nucleus, thereby promoting leaf senescence (Chai et al., 2014). Based on the dose dependent effect of SA, Pasternak et al. (2019) proposes that at low levels it acts as a developmental regulator and at high levels it acts as a stress hormone.

Autophagy is associated with plant senescence as defective mutants display early and strong yellowing leaf symptoms (Hanaoka et al., 2002; Xiong et al., 2005; Avila-Ospina et al., 2014; Li et al., 2014). Autophagy negatively regulates cell death by controlling NPR1-dependent SA signaling during senescence in Arabidopsis (Yoshimoto et al., 2009). The senescence process always accompanies with the equilibrium between oxidative and antioxidative capacities of the plant, which creates a characteristic oxidative environment resulting in the production of reactive oxygen species (ROS) and more toxic derivatives (Bhattacharjee, 2005). Moreover, autophagy is involved in the degradation of oxidized proteins under oxidative stress conditions in Arabidopsis (Xiong et al., 2007). Actually, there is a complicated interplay between ROS and autophagy, i.e., ROS can induce autophagy while autophagy be able to reduce ROS production (Signorelli et al., 2009). Our previous studies showed that LCSA application delays senescence by enhancing the activities of antioxidant enzymes and restricting reactive oxygen species (ROS) accumulation in MeJA-treated leaves (Ji et al., 2016). However, it is still unclear whether autophagy pathway is implicated in the LCSA-alleviated leaf senescence.

Here, the interactions between SA and MeJA in plant senescence were further investigated. By applying transcriptome and interaction network analysis, we identified autophagy-related (ATG) gene modules. In contrast to wild type (Col-0), LCSA cannot alleviate the leaf yellowing phenotype in autophagy defective mutants upon MeJA treatment. Further results revealed that LCSA increased the number of autophagic bodies and autophagic flux during MeJA-induced leaf senescence. Collectively, our work sheds light on up-regulation of autophagy by LCSA is a key regulator to alleviate MeJA-induced leaf senescence.

## MATERIALS AND METHODS

### Plant materials and hormone treatments

Arabidopsis plants of wild-type (WT, Col-0), *atg5-1* (SAIL_129_B07), *atg7-2* (GK-655B06) and eYFP-ATG8e (Zhuang et al., 2013) were grown in a greenhouse at 22 °C with 16 h light photoperiod (120 μmol quanta^−2^ m^−2^). Phytohormones treatment was performed as described by Ji et al (2016). Briefly, the 3rd and 4th rosette leaves from four weeks of plants were detached and incubated in 3 mM MES buffer (pH 5.8) containing 50 μM methyl jasmonate (MeJA) and/or 10 μM salicylic acid (SA). MeJA was prepared from a 50 mM stock solution in ethanol. Solutions without MeJA were supplemented with equal amounts of ethanol.

### Photochemical efficiency and chlorophyll content measurements

The photochemical efficiency was measured with an Imaging-PAM Chlorophyll Fluorometer (PAM-MINI, Walz, Germany) followed the procedure described previously (Zhou et al., 2015). After dark-adapted for 1 h, parameters Fo (minimum fluorescence with PSII reaction centers fully open) and Fm (maximum fluorescence after dark adaptation) were acquired with a 0.8-s saturating pulse (4,000 μmol photons m^-2^ s^-1^). The value of Fv/Fm was calculated by the formulas (Fm-Fo)/Fm. Total Chlorophyll was determined as reported by Coombs et al. (1987). Chlorophyll was extracted by immersion in 90% ethanol at 65 °C for 2 h. The absorbance at 664 nm and 647 nm were determined with a Lambda 35 UV/VIS Spectrometer (Perkin-Elmer) (Zeng et al., 2016). The concentration per fresh weight of leaf tissue was calculated according to the formula: micromoles of chlorophyll per milliliter per gram fresh weight = 7.93(A664) + 19.53(A647). The percentages of Fv/Fm and chlorophyll content are calculated relative to the initial levels of samples before treatment (time zero).

### RNA-Seq analysis

Detached 3rd and 4th rosette leaves from 4-week old plants were immersed in 3 mM MES buffer (pH 5.8) containing 10 μM SA, 50 μM MeJA, and MeJA together with SA for 24 h. Total RNA for RNA-Seq was extracted from leaves using a Hipure plant RNA kit (Magen, China). Purified RNA was analyzed either using a ND-1000 Nanodrop (Thermo Fisher, USA), or by agarose gel electrophoresis to determine the RNA quantity. Those RNA samples with no smear seen on agarose gels, a 260/280 ratio above 2.0, and RNA integrity number greater than 8.0 were used. For RNA-Seq analysis, we mixed three replication samples for each treatment into one, and total RNA samples were then sent to RiboBio Co., Ltd (Guangzhou, China) for sequencing. The NEBNext Poly(A) mRNA Magnetic Isolation Module (NEB, USA) was used for mRNA purification. The Ultra II RNA Library Prep Kit for Illumina was used for RNA library construction. The libraries were sequenced as 50-bp single end reads using Illumina Hiseq2500 according to the manufacturer’s instructions.

### Differential Expression Analysis

Raw read count of each gene was generated using HTSeq with union-count mode (Love et al., 2014). After normalization by Reads Per Kilobase per Million mapped reads (rpkm), normalized read count table was used for determining differentially expressed genes (DEGs) (Anders and Huber, 2010; Cheng et al., 2016a, 2016b), which were defined as those with 2-fold changes. Fold change was calculated using log2 (normalized read count+1). An R package *clusterprofiler* was used to perform the functional category analysis to detect the significantly enriched Gene Ontology (GO) terms (Cheng and Leung, 2018a, and 2018b). Significantly enriched GO terms were selected by a threshold of p ≤ 0.05. Protein-protein interaction (PPI) data was obtained from the STRING database (v.10, http://string-db.org) (Szklarczyk et al., 2014). To construct a high-confidence network, only the PPIs with confidence scores larger than 0.7 were considered in this work. ClusterONE was adopted for the identification of protein clusters or functional modules using default parameters as described previously (Cheng et al., 2017; Cheng et al., 2019). The protein modules including five or more than five members and having connection density over 0.5 are defined as modules.

### RT-qPCR

Total RNA was isolated using Eastep Super RNA Kit (Promega, Shanghai, China) and genomic DNA was removed using DNase I. 1 μg of RNA was used to make cDNA with the GoScript™ Reverse Transcription System (Promega, Shanghai, China). For qPCR 10 μL of Green-2-Go 2X qPCR-S Mastermix (Sangon, Shanghai, China) and 1 μL of cDNA (100 ng/μL) for a total of 20 μL was used in each well. Real-time PCR was done on a CFX Connect Real-Time System (BioRad) at 95 °C for 2 mins, and 45 cycles of 95 °C for 15 s, 55 °C for 30s, and 72 °C for 30s followed by a melting curve analysis. For each sample 3 biological reps were used and repeated 3 times for technical replication. qPCR was analyzed using the ΔΔCt method. Primers for qPCR were showed in Table S2. Statistical significance was determined using Duncan’s multiple range test.

### Confocal microscopy

Detached 3rd and 4th rosette leaves were immersed in 3 mM MES buffer (pH 5.8) containing 50 μM MeJA and/or 10 μM SA for 24 h. Confocal images were captured with 63x (numerical aperture [NA], 1.4) objective using an LSM 880 microscope (Zeiss). For quantification of autophagic puncta, randomly selected 15 to 20 images for each three independent experiments were quantified with ImageJ. All images were collected with the same settings determined prior to the experiment to yield nonsaturating conditions.

### Western Blot

Treatment leaves were ground in liquid nitrogen and extracted with the lysis buffer (50 mM Tris-HCl, pH 7.5, 150 mM NaCl,1 mM EDTA,1% SDS, and 1x Protease Inhibitor Cocktail (MCE, Medchemexpress)). The total lysate was centrifuged at 12,000g for 20 min at 4°C. Protein extracts were separated using SDS-PAGE and then western blotted. Blots were stained with Ponceau S to confirm even loading. A 1:10000 dilution of monoclonal rabbit anti-GFP antibody (HuaAn Biotechnology, Hangzhou, China) was used. Subsequently, blots were washed and incubated with an anti-Rabbit IRDye® 800CW conjugated secondary antibody (Abcam).

## RESULTS

### LCSA delays MeJA-induced leaf senescence

Our previous results indicated that SA delays MeJA-induced leaf senescence in a concentration-dependent manner, showing accelerated by high SA concentrations (greater than 100 μM) but attenuated by low SA concentrations (1-50 μM) (Ji et al., 2016). On this basis, 10 μM SA, the most effective concentration according to Ji et al., 2016, was selected as low working solution to further confirm the effect of LCSA. As shown in Figure 1, in contrast to control, LCSA did not appear to have a discernible effect on senescence. Leaves incubated with MeJA (50 μM) were greatly turned yellow after 5 days treatment. However, when MeJA worked together with LCSA (MeJA+LCSA), the leaf yellowing was alleviated (Figure 1A). Consistent with the visible phenotype, the photochemical efficiency Fv/Fm and loss of chlorophyll content in the leaves combined treatment with LCSA and MeJA was less severe relative to that of the leaves treated with MeJA alone (Figure 1B and 1C). These physiological and biochemical data is consistent with our previous finding that LCSA provide protection against senescence caused by MeJA.

**Figure 1.**
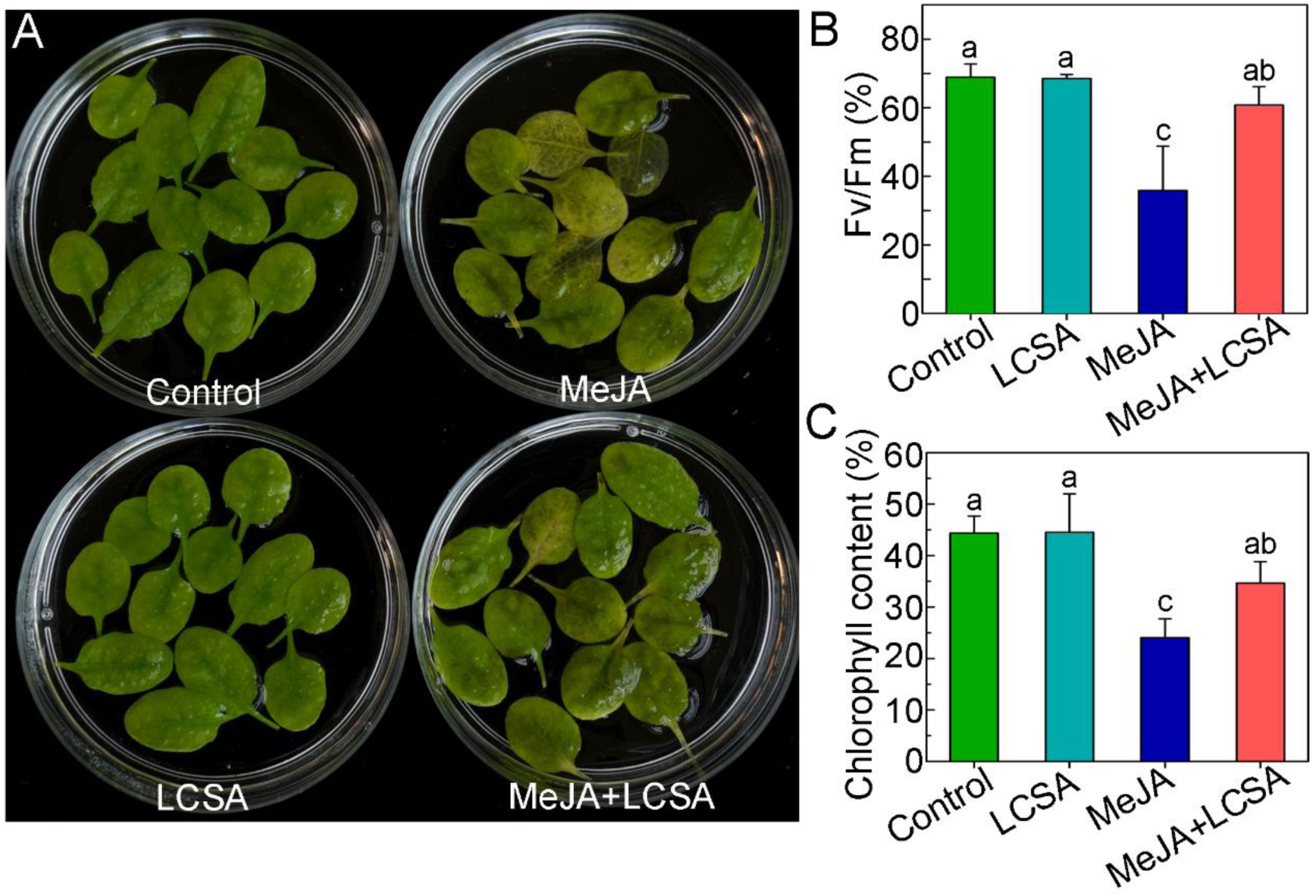
LCSA alleviates MeJA-induced leaf senescence. (A) Phenotypes of detached leaves under LCSA and/or MeJA treatments. The 3rd and 4th rosette leaves were incubated in 3 mM MES buffer (pH 5.8) containing LCSA (10 μM) or MeJA (50 μM) alone or in combination (MeJA+LCSA) under continuous light for 5 d. (B and C) Measurement of the maximum quantum efficiency of photosystem II (PSII) photochemistry (Fv/Fm) (B) and total chlorophyll content (C) after LCSA and/or MeJA treatments. The percentages of Fv/Fm and chlorophyll content are relative to the initial levels at time zero. Data are the mean ± SE of three independent experiments. Different letters indicate statistically significant differences between each treatment (Duncan’s multiple range test, p < 0.05).

### Expression patterns of genes in LCSA-induced delayed leaf senescence

To investigate the genome-wide effect of LCSA on MeJA-induced gene expression changes, we performed RNA-sequencing experiments. Since gene transcription regulation occurs prior to visible phenotype, leaves treatment with phytohormones at 1 d were selected according to our previous study (Ji et al., 2016). Totally, 408, 2536 and 2800 genes displayed at least 2-fold changes in the expression level of LCSA, MeJA, and MeJA+LCSA-treated leaves, respectively, relative to control leaves (Figure 2A). Of these, the number of differentially expressed genes (DEGs) of LCSA alone were greatly less than that in MeJA or MeJA+LCSA treatment group, in consistent with the inconspicuous phenotype between SA and control leaves (Figure 1). Therefore, our study is mainly concentrated on the differential expression of gene between MeJA and MeJA+LCSA.

**Figure 2.**
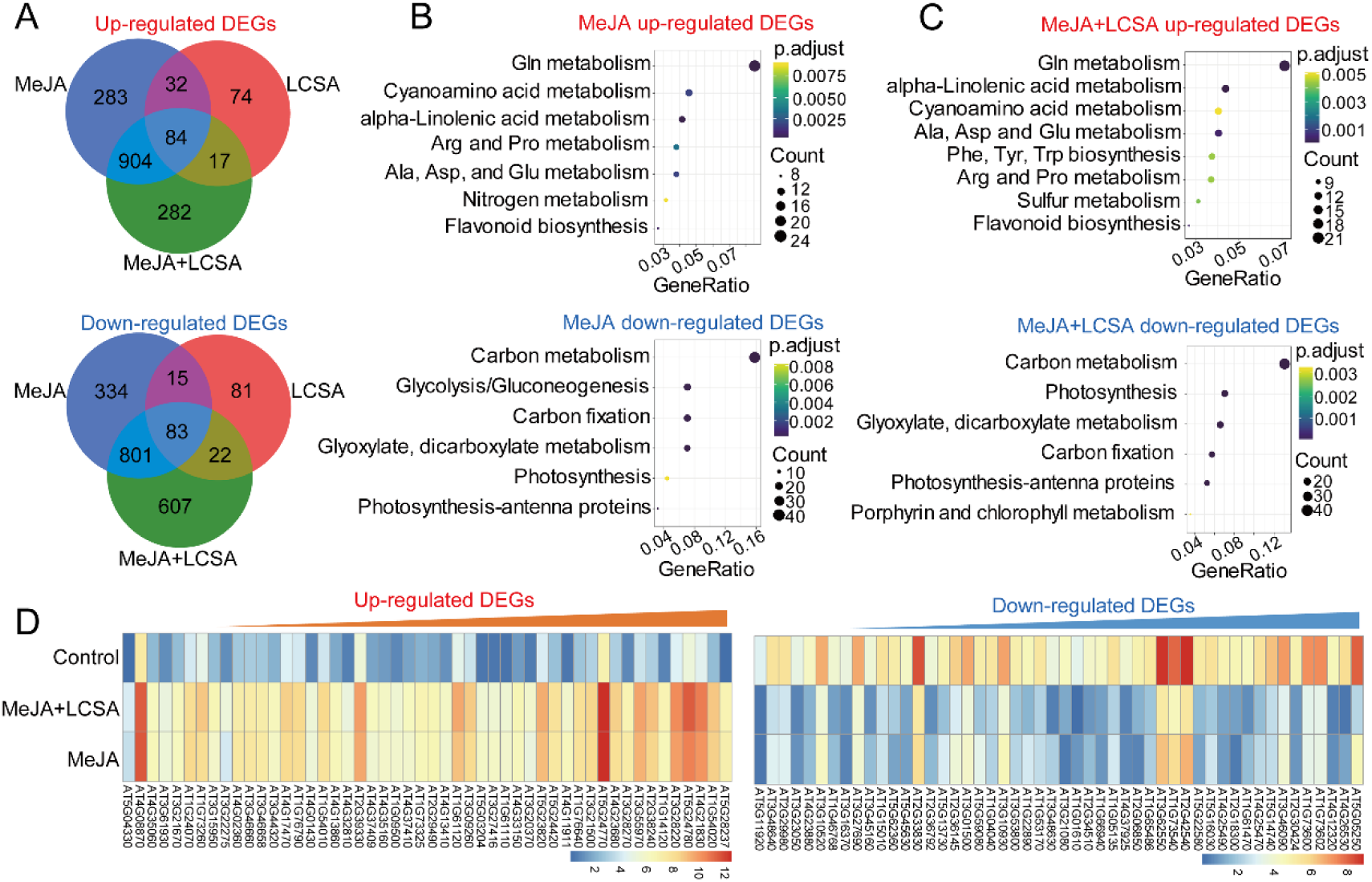
RNA-Seq analyses of differentially expressed genes (DEGs) in samples treated with LCSA, MeJA and LCSA+MeJA. (A) Venn diagram showing the overlap of DEGs between LCSA, MeJA and LCSA+MeJA-treated samples. (B and C) The pathway enrichment analysis of up or down-regulated DEGs induced by MeJA alone (B) or LCSA+MeJA (C). (D) The heatmap showing expression of top 50 up-regulated and down-regulated DEGs between MeJA and MeJA+LCSA treatment group.

To interpret the upregulated and down-regulated DEGs resulting from the MeJA and MeJA+LCSA treatment, functional enrichment of Gene Ontology (GO) terms was performed using the hypergeometric test (P-value < 0.05). The analysis of biological process GO terms illustrated that most of the induced DEGs related to amino acid (Glutathione, Cyanamino acid, arginine, proline, alanine, aspartate and glutamate) metabolism, nitrogen metabolism, and flavonoid biosynthesis, whereas, the repressed DEGs mainly related to carbon metabolism, photosynthesis, and glycolysis (Figure 2B). These features of nitrogen and carbohydrate metabolism are consistent with the senescing phenotype of leaves. In contrast to MeJA alone, unexpectedly, LCSA together with MeJA treatment did not make much differences on the enriched biological processes (Figure 2C). The heatmap illustrated the top 50 up-regulated and down-regulated DEGs, which also revealed an extremely similar expression pattern between the DEGs of MeJA and MeJA+LCSA treatment (Figure 2D). These results indicated that LCSA does not appear to have dominant effects on the genetic regulatory network of basal metabolism like nitrogen metabolism, photosynthesis, and glycolysis.

### Network analysis identifies autophagy-related gene module

Since the enrichment analysis only provided undifferentiated biological processes about basal metabolism, network analysis was conducted using DEGs that induced by MeJA and MeJA+LCSA, respectively. The protein-protein interactions (PPI) were collected from the STRING database, and only the PPIs with confidence scores higher than 0.7 were selected, resulting in a high confidence network with 719964 interactions and 17372 proteins. ClusterONE was used to identify functional protein modules, which were defined by the protein clusters including five or more than five members and having connection density over 0.5 (Cheng et al., 2019). According to such screening specifications, we identified 15 gene modules in MeJA treatment group and 16 gene modules in MeJA+LCSA group, respectively (Figure S1 and S2). Of these, six gene modules were specially detected in the MeJA treatment group (Figure S3). Interestingly, MeJA together with SA exclusively induced seven gene modules, covering genes involved in autophagy-related (ATG) pathway, phytohormone response, ATP-binding cassette transporters, aquaporins, and flavonoid biosynthesis (Figure 3A). In this context, autophagy is an essential intracellular degradation system that plays important roles in nutrient remobilization during leaf senescence (Avila-Ospina et al., 2014). We found that the transcript abundance for ATG proteins (ATG4, ATG8, ATG9, and ATG12) was differentially sensitive to the MeJA+LCSA treatment. From the enriched biological processes and molecular functions, we observed that these ATGs are the core components that contribute to autophagosome mature and biogenesis (Figure 3B and 3C). Collectively, these results suggest a framework in which MeJA together with LCSA regulates the abundance of specific gene network, such as the autophagy process.

**Figure 3.**
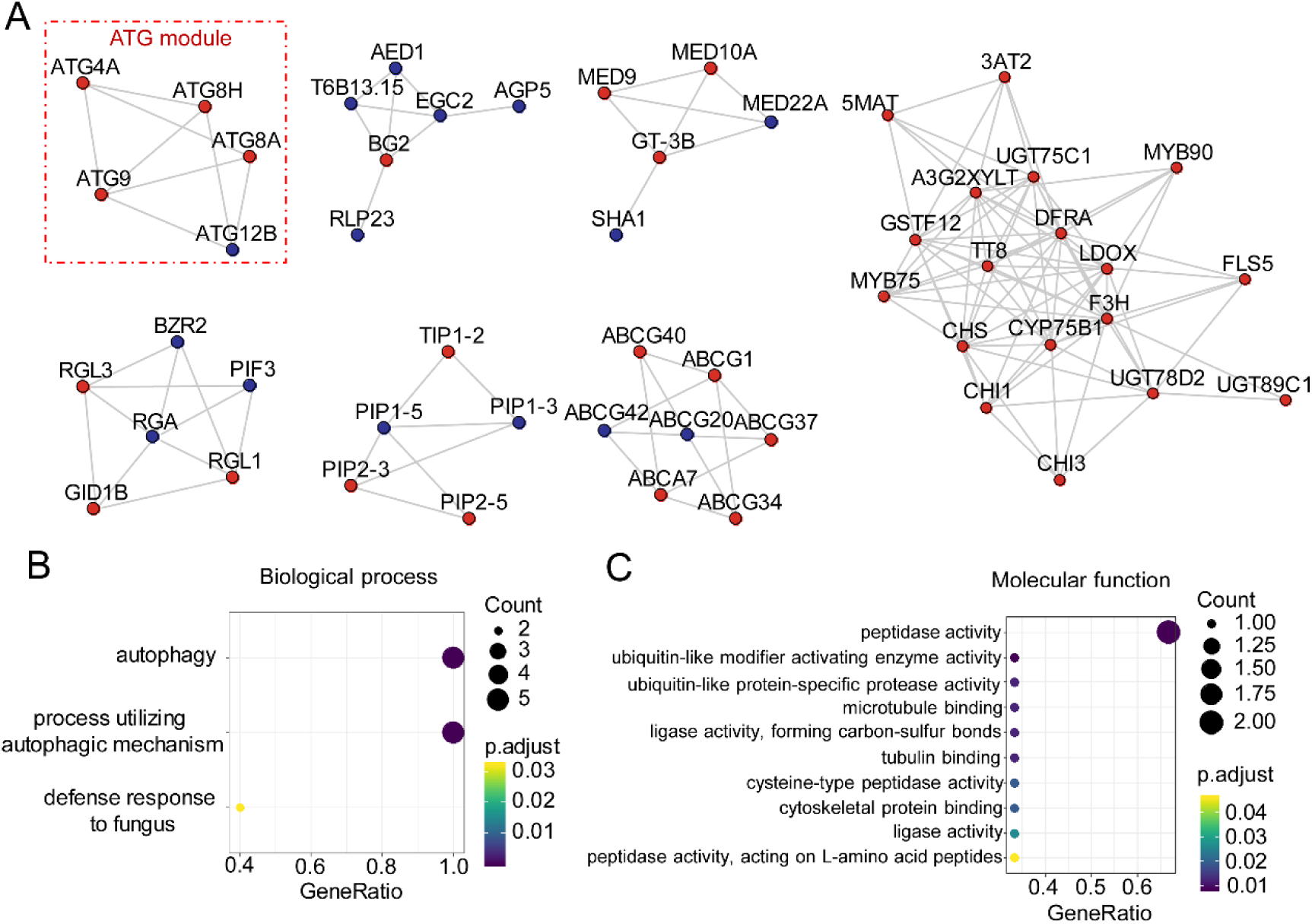
Network analysis identifies distinct signal modules in the DEGs exclusively induced by LCSA+MeJA treatment. (A) Interconnected clusters enriched among the 889 genes and their interactions with neighboring genes. The autophagy specific module was drawn in a red dotted line. Genes are colored in red if they are induced and in blue if they are repressed. (B and C) Biological process (B) and molecular function (C) classification in gene ontology analysis of the DEGs that identified in coexpression networks.

We next investigated whether the autophagy pathway was involved in LCSA-delayed leaf senescence. Ten ATG genes (ATG4A, ATG4B, ATG5, ATG6, ATG7, ATG8A, ATG8E, ATG8H, ATG12A, and ATG12B) that implemented in autophagosome formation were examined by RT-qPCR (Figure 4). In contrast to MeJA alone, most of these determined ATG genes, except for ATG8A and ATG8E, were up-regulated by the combined treatment group (MeJA+LCSA). The differential gene expression of ATG8 isoforms is possible due to they have different expression pattern in distinct tissues (Hanaoka et al., 2002). Interestingly, it should be mentioned that MeJA together with LCSA did not stimulate a much more increase in gene expression compared with control, especially LCSA treatment (Figure 4). These results indicate that restoration of ATG genes expression is closely related to LCSA-delayed leaf senescence.

**Figure 4.**
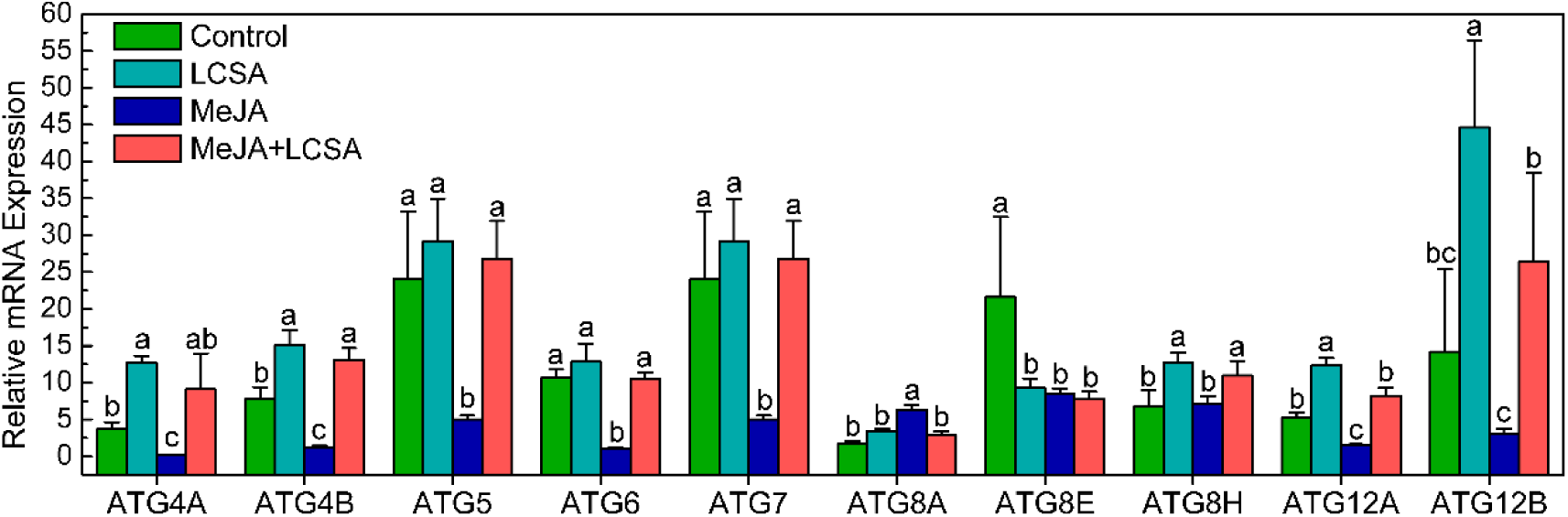
RT-qPCR confirmation of differentially expressed genes that involved in regulation of autophagy. The relative mRNA expression levels were calculated using the ΔΔCt method. The value of each ATG genes were relative to the initial levels at time zero of treatment. Data were the mean ± SE of three independent experiments. Different letters in each genes indicate statistically significant differences between the treatments (Duncan’s multiple range test, p<0.05).

### SA-delayed leaf senescence is dependent on a functional autophagy pathway

To further resolve whether autophagy pathway was crucial for LCSA-delayed leaf senescence, two autophagy defective mutants (*atg5-1* and *atg7-2*), that involved in ATG8 lipidation during phagophore elongation (Feng et al., 2014), were analyzed upon LCSA and/or MeJA treatment. In contrast to wild type (Col-0), leaves from *atg5-1* and *atg7-2* mutants were showed much more yellowing after incubated with MeJA for 5 days (Figure 5A). As expect, the leaf yellowing phenotype was not alleviated when MeJA worked together with LCSA (Figure 5A). Consistently, the photochemical efficiency Fv/Fm in the *atg5-1* and *atg7-2* mutant leaves treated with MeJA+LCSA was not restored relative to that of the leaves treated with MeJA (Figure 5B). Similarly, none of the two mutants had recovered relative chlorophyll content as the Col-0 after combined treatment with MeJA and LCSA (Figure 5C). These genetic results clearly illustrated that the protection against MeJA-induced senescence by LCSA is dependent on a functional autophagy pathway.

**Figure 5.**
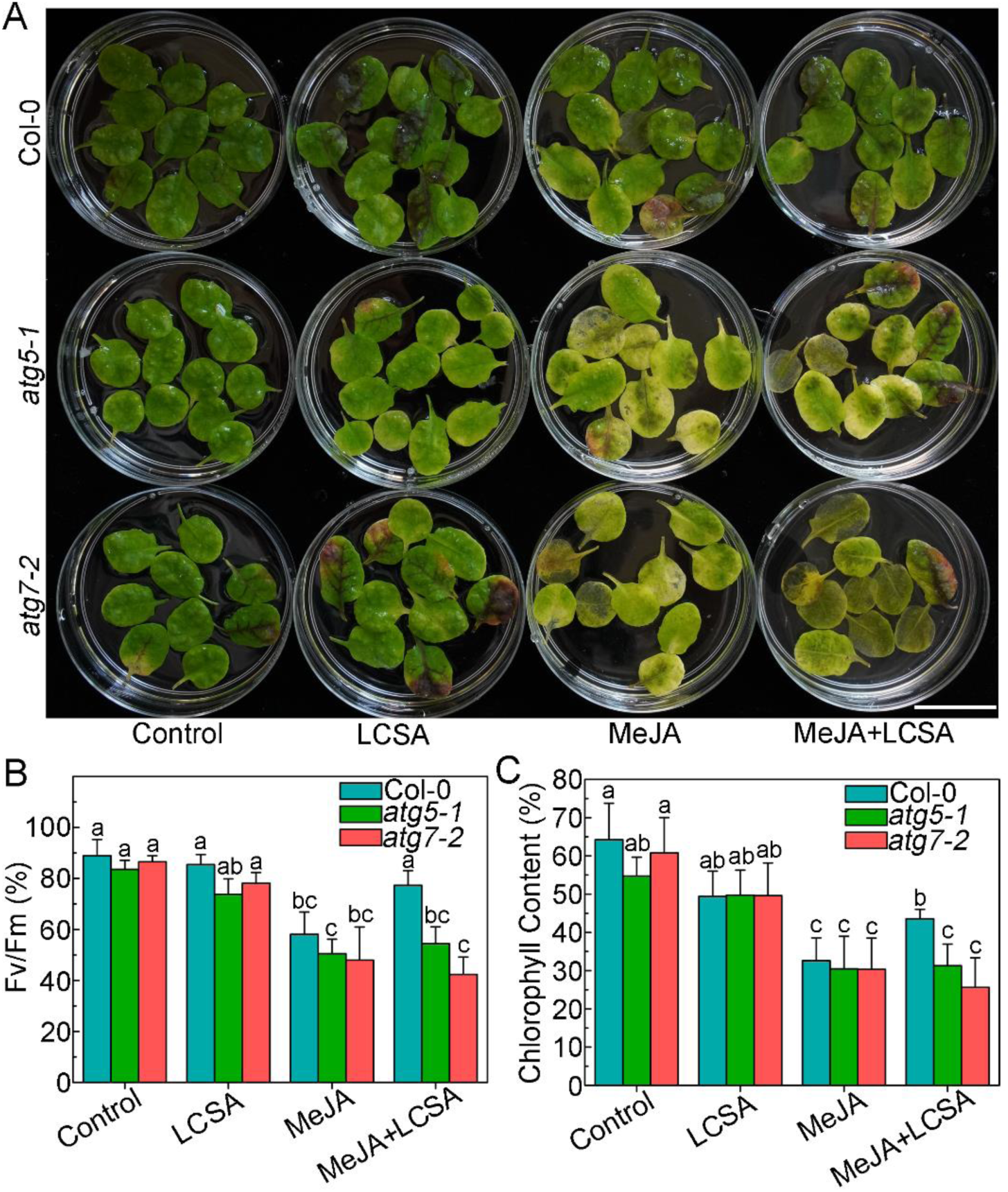
Defective in autophagy restrains the effect of SA on the senescence symptoms. (A) Phenotypes of LCSA-alleviated senescence in Col-0 and autophagy defective mutants (*atg5-1* and *atg7-2*). Detached leaves form four-week-old Col-0, *atg5-1*, and *atg7-2* plants were transferred to MES buffer (pH 5.8) containing LCSA (10 μM) or MeJA (50 μM) or both MeJA and LCSA under continuous light and photographs were taken after 5 days of treatment. (B and C) Relative Fv/Fm (B) chlorophyll levels (C) in the leaves of the Col-0, *atg5-1*, and *atg7-2* described in (A). The data are means ± SD (n = 3) calculated from three biological replicates. Different letters indicate statistically significant differences between the treatments (Duncan’s multiple range test, p<0.05).

### SA increases autophagy activity upon MeJA-induced leaf senescence

Since autophagy pathway was verified involved in LCSA-delayed leaf senescence, we next further determined the detailed autophagy activity. Wild-type Arabidopsis plants expressing the eYFP-ATG8e fusion protein were subjected to LCSA and/or MeJA treatment, and the effects of LCSA on autophagy activity were analyzed by confocal microscopy of the YFP fluorescence. In control and LCSA treatment conditions, we observed a few fluorescent punctate structures that were identified previously as ATG8-tagged autophagosomes (or autophagic bodies) (Yoshimoto et al., 2004; Contento et al., 2005; Thompson et al., 2005). Incubation of MeJA alone induced a slightly increase in accumulation of autophagic bodies (Figure 6A). However, when the detached leaves were subjected to combined treatment with MeJA and LCSA, there was a greatly increase in the fluorescent vesicles (Figure 6A). The statistical results showed that the number of autophagic bodies was more than 2-fold higher in MeJA+LCSA group than that of treatment with MeJA alone (Figure 6B). Western blot analysis also indicated that MeJA together with SA treatment enhanced autophagic flux, showing a decrease in eYFP-ATG8 compared with other treatments (Figure 6C). Taken together, our observations collectively suggest that LCSA activates the autophagy activity to delay MeJA-induced leaf senescence.

**Figure 6.**
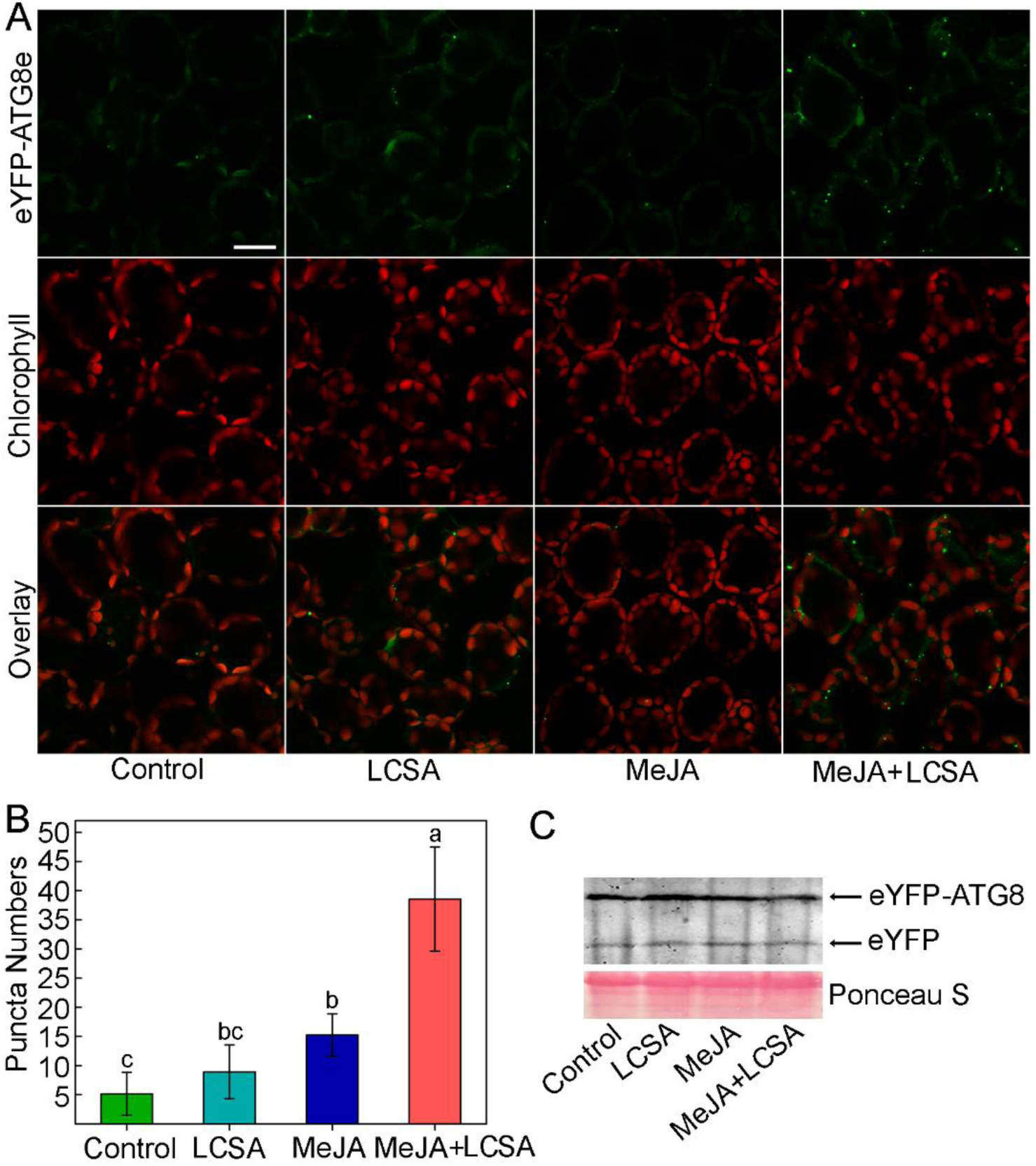
LCSA enhances the formation of autophagosomes upon MeJA-induced leaf senescence. (A) Microscopic analyses of autophagosome-related structures in the eYFP-ATG8e plant under LCSA or MeJA or both MeJA and LCSA treatment. Bar, 20 μm. Statistical analysis of the puncta numbers displayed in (A). The number of puncta was calculated per 0.01 mm^2^ from at least 15 pictures. Different letters indicate statistically significant differences between the treatments (Duncan’s multiple range test, p<0.05). (C) Immunoblot analysis of eYFP-ATG8 processing. Crude extracts were subjected to SDS-PAGE and immunoblot analysis with anti-GFP antibodies. The amount of protein loaded per lane was stained by ponceau S.

## DISCUSSION

As the final stage of leaf development, leaf senescence is a complex process that involves thousands of genes and multiple layers of regulation. Mechanisms governing the specificity regulation of phytohormones and output gene expression are therefore of great interest. The primary objective of the work is to further explore the crosstalk between SA and JA signaling in regulating plant leaf senescence. We have concentrated on examining the mechanisms likely to underpin changes in the transcriptome in response to LCSA and/or MeJA. Specifically, an autophagy module was identified from the DEGs that exclusively induced by MeJA together with SA (Figure 3). Further results demonstrate that the upregulation of autophagy by LCSA serves important function in alleviating MeJA-induced leaf senescence (Figure 5 and 6).

Previously, we found that SA delays MeJA-induced leaf senescence in a concentration dependent manner (Ji et al, 2016). The dosage-dependent effect of SA also has been reported in plant root meristem regulation. SA at low levels (below 50 µM) promotes adventitious roots and alters architecture of the root apical meristem, whereas high-concentration SA (>50 µM) inhibits root growth (Pasternak et al., 2019). Such discrepancies are probably due to SA acts as a developmental regulator at low levels, but acts as a stress hormone at high levels (Pasternak et al., 2019). Interestingly, RNA-Seq results showed that the number of DEGs in LCSA alone treatment were less than MeJA or LCSA and MeJA combined treatment group (Figure 2A), which consistent with LCSA itself did not have a discernible effect on senescence, showing the inconspicuous phenotype between LCSA and control leaves (Figure 1). Moreover, in contrast to MeJA alone, LCSA together with MeJA treatment did not make much differences on the biological process of GO terms (Figure 2C). These results indicated that LCSA at low level is more likely function as a signaling regulator, which does not have a marked impact on the basal metabolism at least at the genetic regulatory level.

Autophagy promotes cell survival by adapting cells to stress conditions both in plants and mammals. Recent reverse-genetic studies have revealed that autophagy is closely associated with plant senescence, and autophagy defective mutants like *atg2, atg5* and *atg7* all showed early yellowing leaf symptoms (Doelling et al., 2002; Yoshimoto et al., 2009). SA is one of the most promising phytohormones that contribute to the induction of autophagy under stress. It has previously been reported that autophagy negatively regulates cell death by controlling NPR1-dependent SA signaling during senescence in Arabidopsis (Yoshimoto et al., 2009). Here, the ClusterONE was applied to identify discrete gene modules based on PPI network. We identified several modules including autophagy-related network in DEGs that exclusively induced by MeJA together with LCSA (Figure 3A). Importantly, the protection against MeJA-induced senescence by LCSA was abolished in autophagy defective mutant *atg5-1* and *atg7-2* (Figure 5). These data strongly suggest an important role for autophagy in LCSA-alleviated leaf senescence. Notably, unlike the greatly increase of autophagic bodies induced by MeJA+LCSA, autophagosomes under LCSA alone treatment were not statistically significant when compared with control (Figure 6). Nevertheless, it is worth pointing out that SA at 100 µM, a high-concentration that could promote leaf senescence based on our previous study (Chai et al., 2016), greatly induced autophagic structures formation (Figure S4). In this context, we speculate that LCSA might be function like a priming regulator, which could initiate signal amplification and lead to a robust activation of stress response upon MeJA treatment. Actually, the priming induced by some plant activators (e.g. *β*-aminobutyric acid, and thiamine) are dependent on SA signaling (Ahn et al., 2005; Jung et al., 2009; Zhou et al., 2013). It would be interesting to test the priming effect of LCSA on leaf senescence in future research.

In summary, this study further investigated the interactions between SA and MeJA in plant senescence. Several modules including an autophagy-related (ATG) cluster were identified by analyzing the transcriptome data and protein interaction networks. Our results showed that LCSA could upregulate autophagy to alleviate leaf senescence when combined treatment with MeJA. This was confirmed by founding that LCSA cannot alleviate the leaf yellowing phenotype in autophagy defective mutants upon MeJA treatment. Collectively, our work reveals LCSA tend to function as a signaling regulator to upregulate autophagy pathway, which serves as an important cellular mechanism responsible for alleviation of MeJA-induced leaf senescence.

## Data availability

RNA-seq data were deposited in the Sequence Read Archive (SRA) database https://www.ncbi.nlm.nih.gov/sra with accession no. PRJNA578602.

## AUTHOR CONTRIBUTIONS

JZ and LC designed the research. RY, JY, and YJ conducted the experiments. RY, JL, JZ and LC analyzed data. JZ and LC wrote the manuscript. All authors read and approved the manuscript.

## ACKNOWLEDGEMENTS

Thanks for Professor Liwen Jiang (the Chinese University of Hong Kong) for giving the Arabidopsis seeds materials *atg5-1, atg7-2* and eYFP-ATG8e. Thanks for Yang Lv (Fengyuan biotechnology co. LTD, Shanghai, China) for the valuable suggestions for this manuscript. This work was supported by National Science Foundation of China (NSFC) (31600288); Guangdong Provincial Science and Technology Project (2016A020210127); SCNU Youth Teacher Research and Development Fund Project (671075); Scientific Research Projects of Guangzhou (201805010002).

## SUPPLEMENTARY MATERIALS

**Figure S1.**
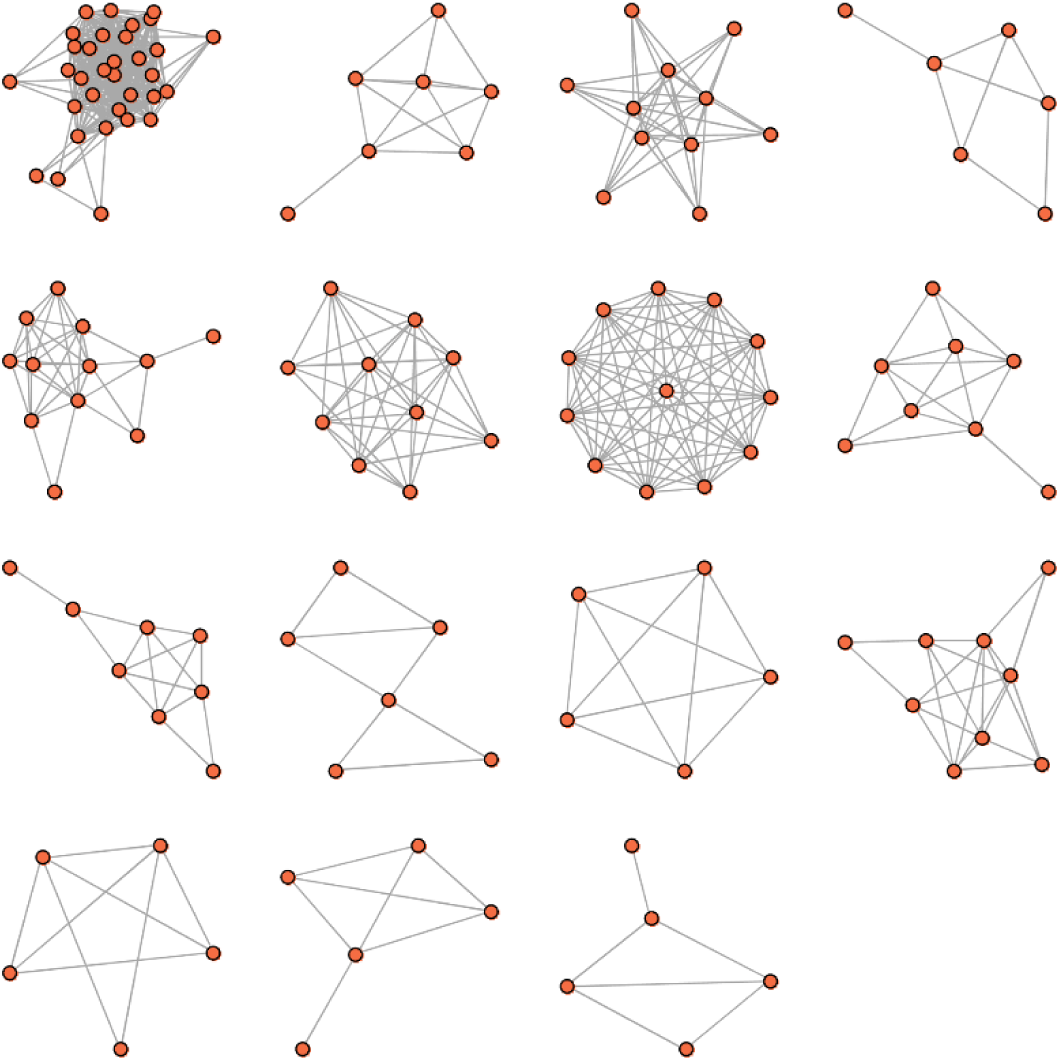
Network analysis identifies gene modules in the DEGs induced by MeJA. Protein modules were identified using ClusterONE, a cluster screen method considering the overlapping neighbor extension. The protein modules including five or more than five members and having connection density over 0.5 are defined as modules.

**Figure S2.**
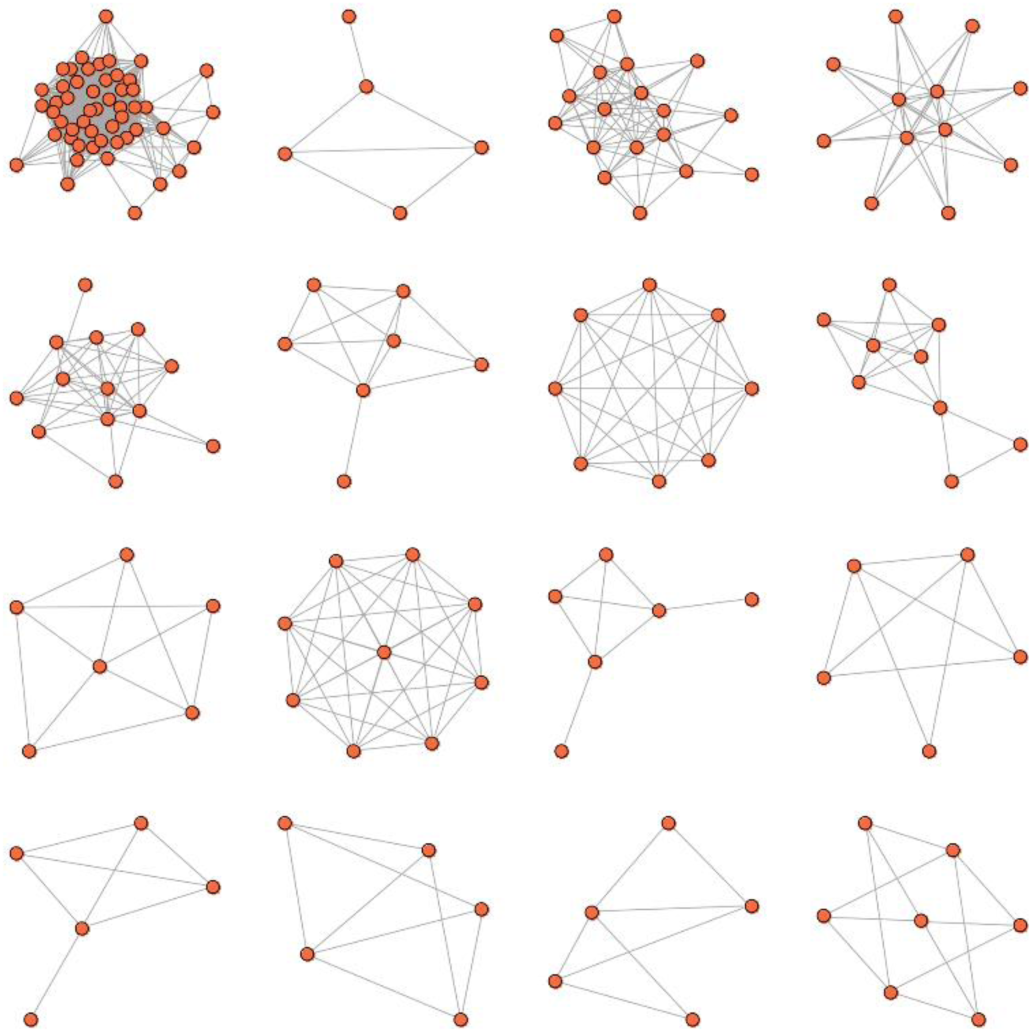
Network analysis identifies gene modules in the DEGs induced by MeJA together with LCSA. Protein modules were identified using ClusterONE. The protein modules including at least five members and having connection density over 0.5 are defined as modules.

**Figure S3.**
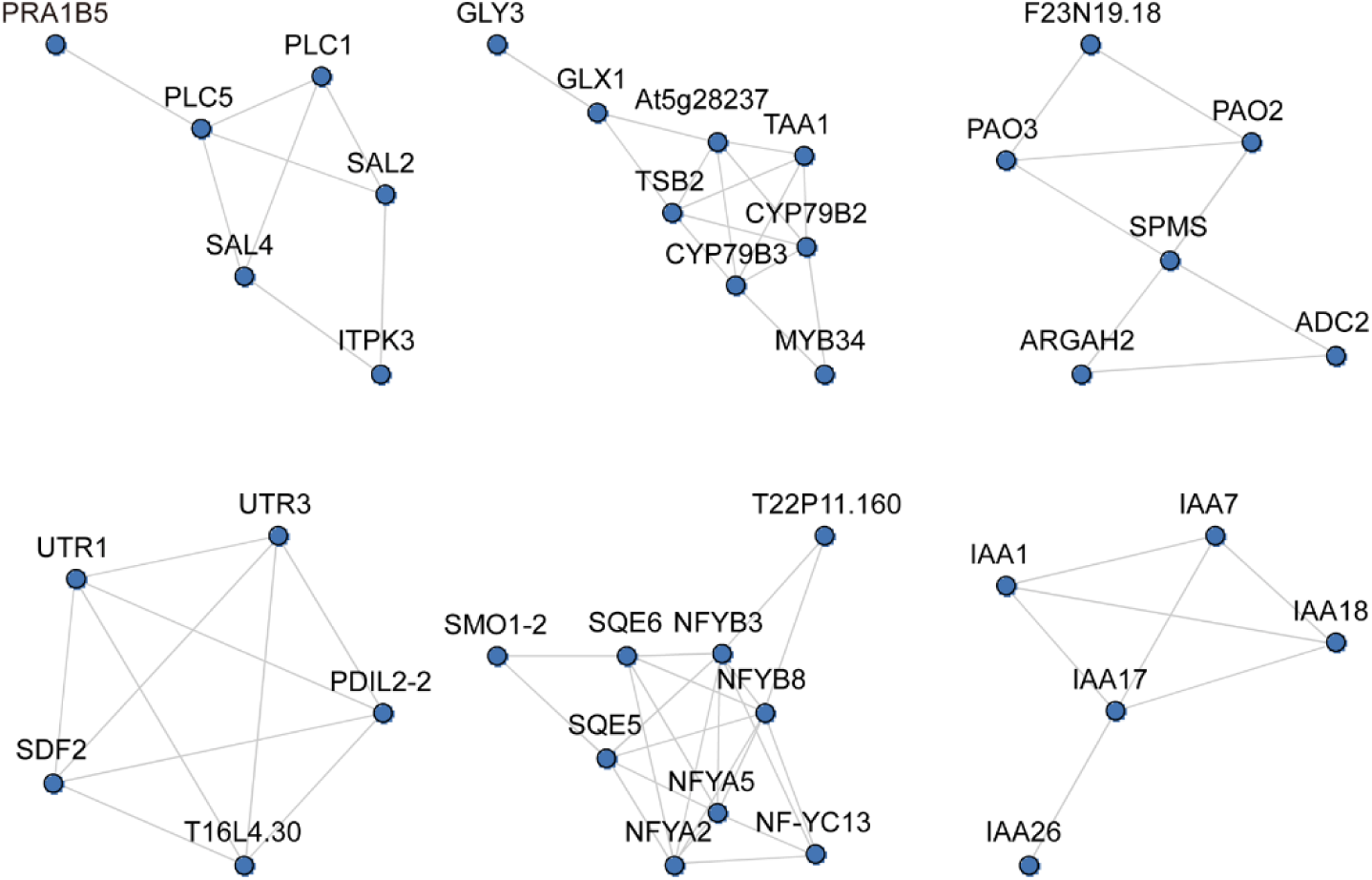
Gene modules exclusively induced by MeJA. After removal of the same modules in MeJA+LCSA treatment group, six gene modules specially induced by MeJA were obtained.

**Figure S4.**
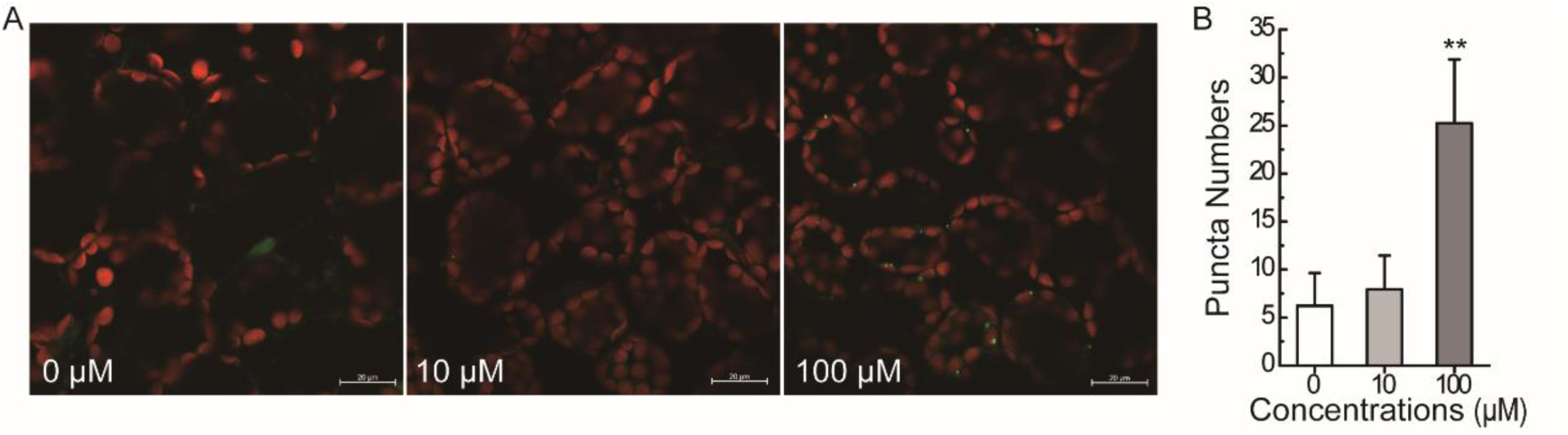
Effect of different concentrations of SA on autophagic puncta induction. Microscopic analyses of autophagic structures in the eYFP-ATG8e plant under 0, 10 and 100 M SA treatment. Bar, 20 μm. (B) Statistical analysis of the puncta numbers displayed in (A). The number of puncta was calculated per 0.01 mm^2^ from at least 15 pictures. Asterisks indicate a significant difference according to Student’s t-test, **P<0.01.

**Table S1.**
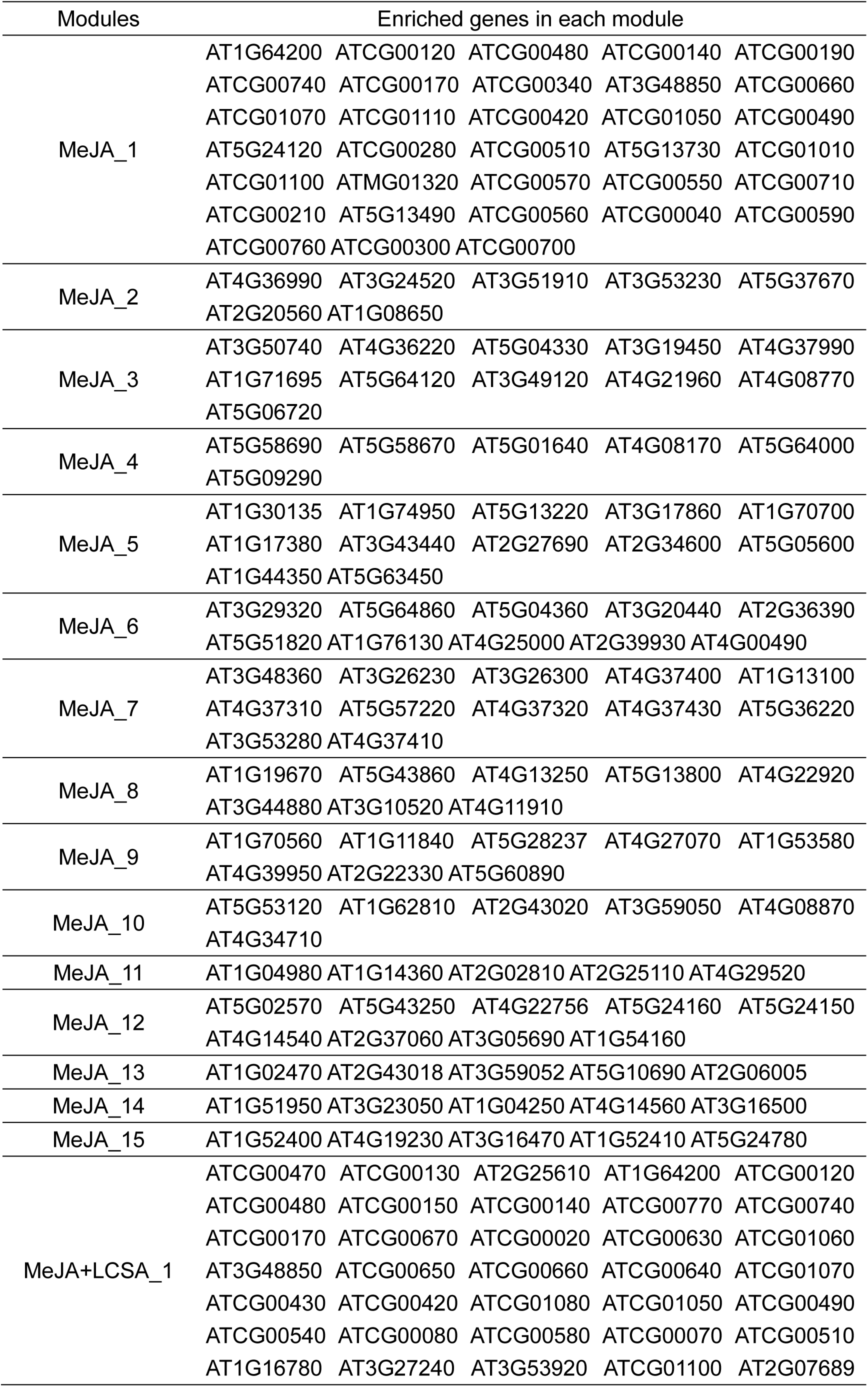
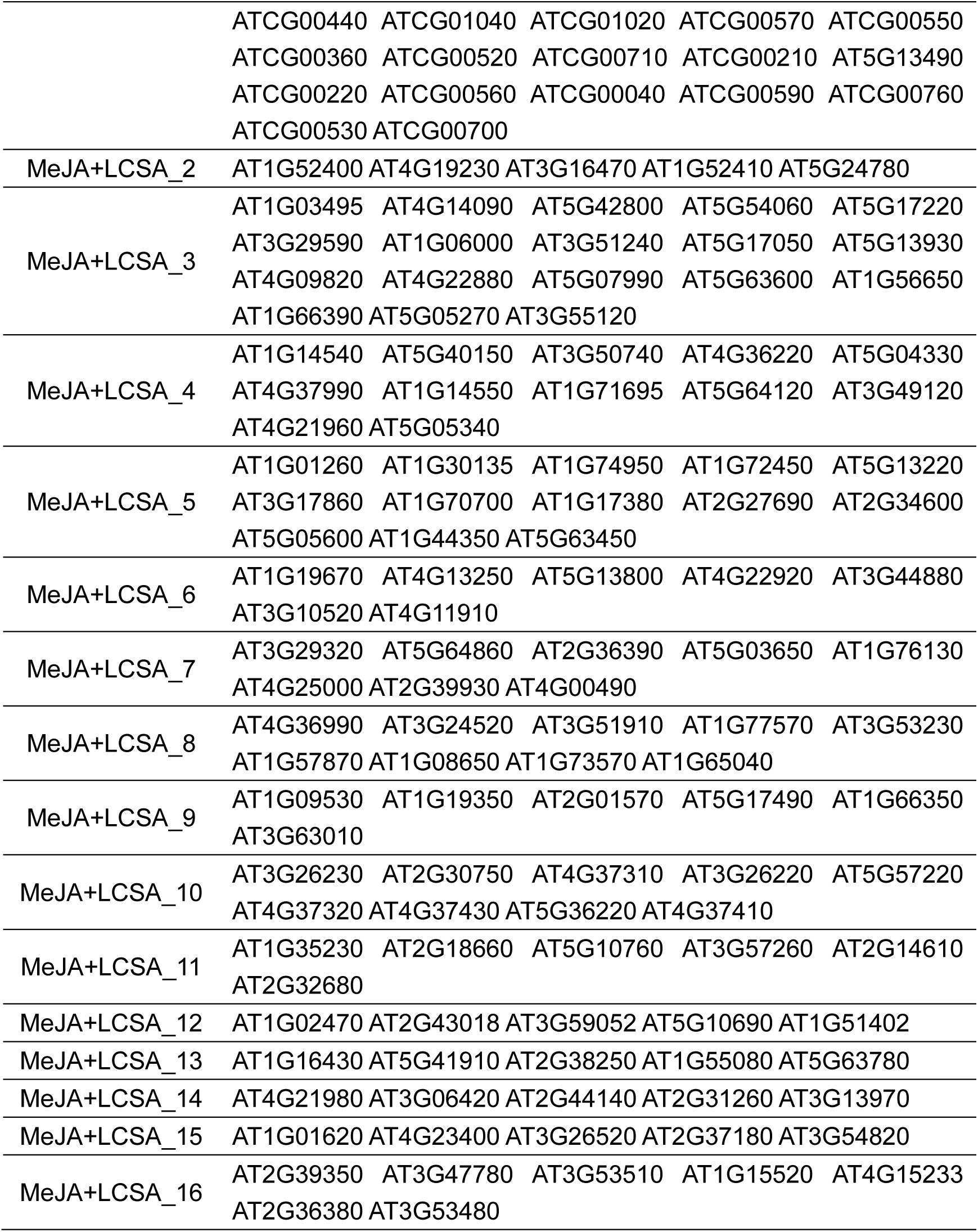
Genes list for enriched modules in the DEGs induced by MeJA and MeJA together with LCSA.

| Modules | Enriched genes in each module |  |  |  |  |
| --- | --- | --- | --- | --- | --- |
| MeJA_1 | AT1G64200 | ATCG00120 | ATCG00480 | ATCG00140 | ATCG00190 |
|  | ATCG00740 | ATCG00170 | ATCG00340 | AT3G48850 | ATCG00660 |
|  | ATCG01070 | ATCG01110 | ATCG00420 | ATCG01050 | ATCG00490 |
|  | AT5G24120 | ATCG00280 | ATCG00510 | AT5G13730 | ATCG01010 |
|  | ATCG01100 | ATMG01320 | ATCG00570 | ATCG00550 | ATCG00710 |
|  | ATCG00210 | AT5G13490 | ATCG00560 | ATCG00040 | ATCG00590 |
|  | ATCG00760 | ATCG00300 | ATCG00700 |  |  |
| MeJA_2 | AT4G36990 | AT3G24520 | AT3G51910 | AT3G53230 | AT5G37670 |
|  | AT2G20560 | AT1G08650 |  |  |  |
| MeJA_3 | AT3G50740 | AT4G36220 | AT5G04330 | AT3G19450 | AT4G37990 |
|  | AT1G71695 | AT5G64120 | AT3G49120 | AT4G21960 | AT4G08770 |
|  | AT5G06720 |  |  |  |  |
| MeJA_4 | AT5G58690 | AT5G58670 | AT5G01640 | AT4G08170 | AT5G64000 |
|  | AT5G09290 |  |  |  |  |
| MeJA_5 | AT1G30135 | AT1G74950 | AT5G13220 | AT3G17860 | AT1G70700 |
|  | AT1G17380 | AT3G43440 | AT2G27690 | AT2G34600 | AT5G05600 |
|  | AT1G44350 | AT5G63450 |  |  |  |
| MeJA_6 | AT3G29320 | AT5G64860 | AT5G04360 | AT3G20440 | AT2G36390 |
|  | AT5G51820 | AT1G76130 | AT4G25000 | AT2G39930 | AT4G00490 |
| MeJA_7 | AT3G48360 | AT3G26230 | AT3G26300 | AT4G37400 | AT1G13100 |
|  | AT4G37310 | AT5G57220 | AT4G37320 | AT4G37430 | AT5G36220 |
|  | AT3G53280 | AT4G37410 |  |  |  |
| MeJA_8 | AT1G19670 | AT5G43860 | AT4G13250 | AT5G13800 | AT4G22920 |
|  | AT3G44880 | AT3G10520 | AT4G11910 |  |  |
| MeJA_9 | AT1G70560 | AT1G11840 | AT5G28237 | AT4G27070 | AT1G53580 |
|  | AT4G39950 | AT2G22330 | AT5G60890 |  |  |
| MeJA_10 | AT5G53120 | AT1G62810 | AT2G43020 | AT3G59050 | AT4G08870 |
|  | AT4G34710 |  |  |  |  |
| MeJA_11 | AT1G04980 | AT1G14360 | AT2G02810 | AT2G25110 | AT4G29520 |
| MeJA_12 | AT5G02570 | AT5G43250 | AT4G22756 | AT5G24160 | AT5G24150 |
|  | AT4G14540 | AT2G37060 | AT3G05690 | AT1G54160 |  |
| MeJA_13 | AT1G02470 | AT2G43018 | AT3G59052 | AT5G10690 | AT2G06005 |
| MeJA_14 | AT1G51950 | AT3G23050 | AT1G04250 | AT4G14560 | AT3G16500 |
| MeJA_15 | AT1G52400 | AT4G19230 | AT3G16470 | AT1G52410 | AT5G24780 |
| MeJA+LCSA_1 | ATCG00470 | ATCG00130 | AT2G25610 | AT1G64200 | ATCG00120 |
|  | ATCG00480 | ATCG00150 | ATCG00140 | ATCG00770 | ATCG00740 |
|  | ATCG00170 | ATCG00670 | ATCG00020 | ATCG00630 | ATCG01060 |
|  | AT3G48850 | ATCG00650 | ATCG00660 | ATCG00640 | ATCG01070 |
|  | ATCG00430 | ATCG00420 | ATCG01080 | ATCG01050 | ATCG00490 |
|  | ATCG00540 | ATCG00080 | ATCG00580 | ATCG00070 | ATCG00510 |
|  | AT1G16780 | AT3G27240 | AT3G53920 | ATCG01100 | AT2G07689 |

|  |  |
| --- | --- |
|  | ATCG00440 ATCG01040 ATCG01020 ATCG00570 ATCG00550<br>ATCG00360 ATCG00520 ATCG00710 ATCG00210 AT5G13490<br>ATCG00220 ATCG00560 ATCG00040 ATCG00590 ATCG00760<br>ATCG00530 ATCG00700 |
| MeJA+LCSA_2 | AT1G52400 AT4G19230 AT3G16470 AT1G52410 AT5G24780 |
| MeJA+LCSA_3 | AT1G03495 AT4G14090 AT5G42800 AT5G54060 AT5G17220<br>AT3G29590 AT1G06000 AT3G51240 AT5G17050 AT5G13930<br>AT4G09820 AT4G22880 AT5G07990 AT5G63600 AT1G56650<br>AT1G66390 AT5G05270 AT3G55120 |
| MeJA+LCSA_4 | AT1G14540 AT5G40150 AT3G50740 AT4G36220 AT5G04330<br>AT4G37990 AT1G14550 AT1G71695 AT5G64120 AT3G49120<br>AT4G21960 AT5G05340 |
| MeJA+LCSA_5 | AT1G01260 AT1G30135 AT1G74950 AT1G72450 AT5G13220<br>AT3G17860 AT1G70700 AT1G17380 AT2G27690 AT2G34600<br>AT5G05600 AT1G44350 AT5G63450 |
| MeJA+LCSA_6 | AT1G19670 AT4G13250 AT5G13800 AT4G22920 AT3G44880<br>AT3G10520 AT4G11910 |
| MeJA+LCSA_7 | AT3G29320 AT5G64860 AT2G36390 AT5G03650 AT1G76130<br>AT4G25000 AT2G39930 AT4G00490 |
| MeJA+LCSA_8 | AT4G36990 AT3G24520 AT3G51910 AT1G77570 AT3G53230<br>AT1G57870 AT1G08650 AT1G73570 AT1G65040 |
| MeJA+LCSA_9 | AT1G09530 AT1G19350 AT2G01570 AT5G17490 AT1G66350<br>AT3G63010 |
| MeJA+LCSA_10 | AT3G26230 AT2G30750 AT4G37310 AT3G26220 AT5G57220<br>AT4G37320 AT4G37430 AT5G36220 AT4G37410 |
| MeJA+LCSA_11 | AT1G35230 AT2G18660 AT5G10760 AT3G57260 AT2G14610<br>AT2G32680 |
| MeJA+LCSA_12 | AT1G02470 AT2G43018 AT3G59052 AT5G10690 AT1G51402 |
| MeJA+LCSA_13 | AT1G16430 AT5G41910 AT2G38250 AT1G55080 AT5G63780 |
| MeJA+LCSA_14 | AT4G21980 AT3G06420 AT2G44140 AT2G31260 AT3G13970 |
| MeJA+LCSA_15 | AT1G01620 AT4G23400 AT3G26520 AT2G37180 AT3G54820 |
| MeJA+LCSA_16 | AT2G39350 AT3G47780 AT3G53510 AT1G15520 AT4G15233<br>AT2G36380 AT3G53480 |

**Table S2.**
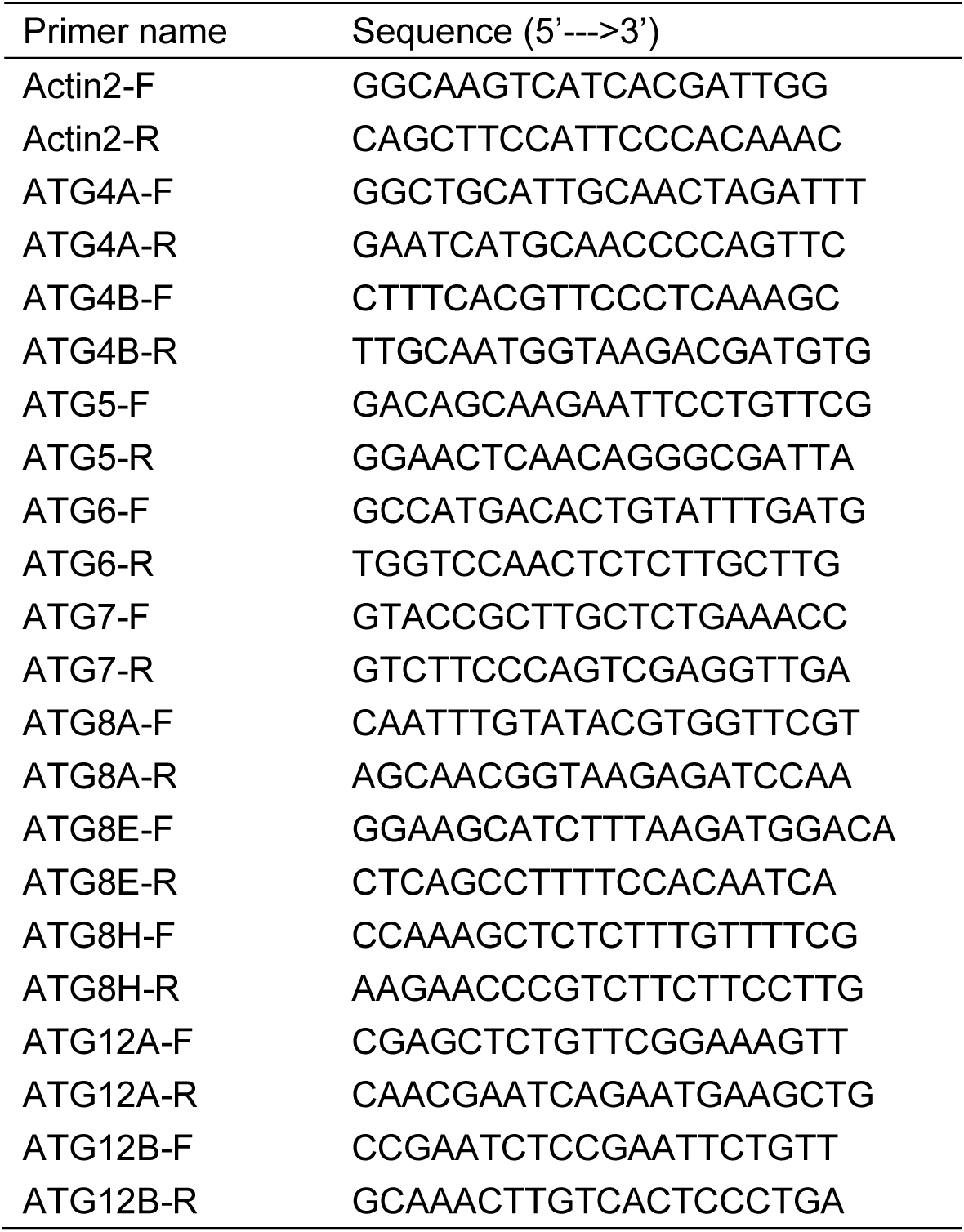
Primers used for RT-qPCR. F, Forward; R, Reverse.

